# Unravelling the Impact of miR-21 Overexpression on the MicroRNA Network and Cancer Pathways

**DOI:** 10.1101/2024.05.10.593469

**Authors:** Meredith Hill, Sarah Stapleton, Phuong Thao Nguyen, Dayna Sais, Fiona Deutsch, Valarie Gay, Deborah Marsh, Nham Tran

## Abstract

MicroRNAs (miRNA, miRs) are small noncoding RNAs that are ubiquitously expressed in all mammalian cells. Their primary function is the regulation of nascent RNA transcripts by direct binding to regions on the target. There is now exciting data to suggest that these miRNAs can bind to other miRNAs, and this may have a broader impact on gene regulation in disease states. The oncomiR miR-21 is one of the highest-expressing miRNAs in cancer cells, and in this study, we characterise which miRNAs could be potential targets of miR-21. In cancer cells delivered with a miR-21 mimic, there was an observable shift of the miRNA milieu. We demonstrate that the miR-17-92a cluster, which harbours six miRNA members, may be a target of miR-21 regulation. Additionally, the primary transcript of miR-17-92a was reduced in the presence of miR-21. In the broader context of gene regulation, overexpression of miR-21 shifted the expression of more than 150 miRNAs, including those known to regulate genes encoding proteins in cancer pathways such as the MAPK signalling and FoxO pathways. This study expands upon our limited understanding of miR:miR regulatory network and reinforces the concept that miRNAs can regulate each other, thereby influencing broader gene regulatory networks.

## Introduction

Gene regulation is a fundamental process in all mammalian cells. Small non-coding RNAs such as microRNAs (miRNAs) are potent gene regulators that can have a profound impact on cell fate. These miRNAs are typically 21-22 nucleotides in length but are derived from much longer RNA transcripts (Lagos-Quintana et al., 2001; Lau et al., 2001; Lee and Ambros, 2001). Moreover, microRNAs act by binding to specific regions on target genes, mostly at 3’UTRs, via their miRNA recognition elements (MRE) sites. Upon binding, microRNAs can then suppress these targets by preventing translation of the RNA message or degradation of the transcript. This results in the suppression of the target gene (Tran and Hutvagner, 2013).

It comes as no surprise that the dysregulation of miRNAs is associated with complex human diseases. We now understand that the aberrant expression of specific miRNAs is linked to multiple cancers of the blood (Calin et al., 2002), breast (Iorio et al., 2005), lung (Eder and Scherr, 2005; Takamizawa et al., 2004), and areas of the head and neck region (Tran et al., 2007). Disease progression is further linked to the temporal nature of miRNA expression and its regulatory impact on multiple pathways. At present, there are more than 2000 human microRNAs recorded in miRbase (Griffiths-Jones, 2006), among which miR-21 stands out as the principal oncogenic miRNA (oncomiRs). This means that miR-21 is often found at elevated levels in cancer cells, where it targets mRNA for genes such as Phosphatase and Tensin Homolog (PTEN) (Meng et al., 2007), Programmed Cell Death 4 (PDCD4) (Lu et al., 2008) and Tropomyosin 1 (TPM1) (Zhu et al., 2007), which have important roles in the pathways leading to tumorigenesis.

The prevailing view is that miR-21 and similar miRNAs play a crucial role in disease progression by binding to and influencing thousands of nascent messenger RNAs. This intricate network relies on the precise expression of these miRNAs. We and others have suggested the possibility of miRNAs also regulating other miRNAs, a concept known as miRNA:miRNA regulation (Hill and Tran, 2021b). This type of interaction was initially identified in Drosophila with certain miRNAs, such as miR-5 and miR-6, as well as miR-9 and miR-79, showing a higher binding affinity for each other than for their intended targets (Lai et al., 2004).

Another case of complementary binding involves miR-107 and let-7, leading to a reduction in mature let-7 levels. The duplex formed by these two mature miRNAs created a series of bulges within its structure, of which the internal loop was vital for the interaction (Chen et al., 2011). However, this interaction raises questions as to how two mature miRNAs may undergo binding whilst bound within the RNA-Induced Silencing Complex (RISC).

Further studies have revealed that miRNAs can mediate direct interactions between a mature miRNA and a primary miRNA in the nucleus. For instance, in murine cardiomyocytes, it was found that the sequence of pri-miR-484 contains an MRE for miR-361 (Wang et al., 2014). The binding of miR-361 to its MRE prevented pri-miR- 484 cleavage by Drosha within the nucleus, which then impacted cardiomyocyte apoptosis.

Additionally, a study revealed that miR-122, predominantly expressed in the liver, can regulate the expression of miR-21 by regulating the expression of the miR-21 primary transcript. The MRE on the pri-miR-21 transcript that is targeted by miR-122 is also recognised by Drosha. Consequently, when miR-122 binds to pri-miR-21, it prevents the cleavage and processing by Drosha, leading to a decrease in the level of mature miR-21 within the cell (Wang et al., 2018). This mechanism has significant implications for cell growth and proliferation, as miR-21 is a known regulator of the tumour suppressor PDCD4 (Lu et al., 2008).

With the emerging concept that miRNAs can influence other miRNAs, we sought to understand the impact of the leading oncomiR, miR-21, on the miRNA milieu in specific cancer cells. We have identified a network of upregulated and downregulated miRNAs as a result of miR-21 overexpression and show evidence that the miR-17-92a cluster is a direct target for miR-21. Furthermore, we demonstrate that overall changes in the miRNA milieu due to miR-21 may impact cancer-specific pathways.

## Methods

### Tissue Culture and miRNA Transfection

The Hypopharyngeal Squamous Cell Carcinoma cell line, UMSCC22B, the Tongue Squamous Cell Carcinoma cell line, SCC4, and HeLa cells were grown in Dulbecco’s Modified Eagle Medium (DMEM) supplemented with 10% Foetal Calf Serum and maintained at 37℃ and 5% CO_2_. At 80-90% confluency, cells were seeded at a density of 5x10^4^ cells per well into a 12-well plate. Once at 60-70% confluency, cells were transfected in triplicate with 10pmol hsa-miR-21-5p mimic (Applied Biosystems^TM^, Thermo Fisher Scientific, Australia), anti-miR-21-5p (Applied Biosystems^TM^, Thermo Fisher Scientific, Australia), scramble/non-targeting probe, and non-transfected cells included as controls. The cells were harvested 24- and 48- hours post-transfection for RNA isolation.

### RNA isolation

RNA isolation was performed using RNAzol RT (MRC, USA) with 250uL of the reagent added per well of the 12-well plate. Samples underwent 4-Bromoanisole (BAN) (MRC, USA) purification before undergoing isopropanol reprecipitation. The RNA was resuspended in DNase/RNase-Free Water, and concentration was measured using the NanoDrop One UV-Vis Spectrophotometer. Samples with a 260/280 and 260/230 ratio between 1.7 to 2.1 were used for subsequent analyses.

### Complementary DNA Synthesis and qPCR

Standardised RNA underwent complementary DNA (cDNA) synthesis for miRNAs using the High-Capacity cDNA kit (Applied Biosystems^TM^, Thermo Fisher Scientific, Australia) The cDNA product was measured using the NanoDrop One UV-Vis Spectrophotometer and the concentration was standardised across samples.

Quantitative PCR (qPCR) was performed for each sample in triplicate using the Step One Plus PCR System or the QuantStudio 12K Flex Real-Time PCR System using the TaqMan Gene Expression Assay (Applied Biosystems, Thermo Fisher Scientific, USA), with a reaction volume of 5uL. The results for each triplicate were analysed using LinRegPCR (Ramakers et al., 2003; Ruijter et al., 2009) and subsequent calculations were performed.

### TaqMan Array and Analysis

Representative UMSCC22B samples transfected with 10pmol of miR-21 and a non- transfected control were nominated for miRNA array analysis. cDNA synthesis was performed using the Megaplex^TM^ Primer Pools, Human Pools Set v3.0 kit (Applied Biosystems^TM^, Thermo Fisher Scientific, USA) and the TaqMan^TM^ MicroRNA Reverse Transcription Kit (Applied Biosystems^TM^, Thermo Fisher Scientific, USA). The TaqMan PreAmp Master Mix (Applied Biosystems^TM^, Thermo Fisher Scientific, USA) was used to amplify the generated cDNA. The cDNA was applied to the TaqMan^TM^ OpenArray^TM^ Human MicroRNA Panel, QuantStudio^TM^ 12K Flex (Applied Biosystems^TM^, Thermo Fisher Scientific, USA) and RT-qPCR was performed using the QuantStudio^TM^ 12K Flex Real-Time PCR System following the manufacturer’s instructions.

Several filtering steps were applied to the output of the OpenArray to identify key miRNAs of interest. Each samples’ results were sorted by target name and combined into one sheet for ease of analysis. Delta-delta-Cq Fold-change analysis was applied in relation to the untransfected sample, with U6snRNA as the reference gene (Livak and Schmittgen, 2001). Amp score greater than >1.1 and Cq confidence greater than >0.8 were used to exclude miRNAs of poor amplification quality. The top-ten most upregulated and downregulated miRNAs were then identified from the remaining list of miRNAs.

### Small RNA Next Generation Sequencing (NGS) and Analysis

Duplicate UMSCC22B samples transfected over 48hrs with 10pmol miR-21, anti- miR-21, and a scramble control were chosen for small RNA NGS. In order to be considered for NGS, samples were required to have a minimum of 100ng total RNA, a 260:280 ratio >2 and a 260:230 ratio >1.7. Small RNA NGS was outsourced to the Ramaciotti Centre for Genomics at the University of New South Wales, Kensington, which included quality control, library preparation and single-end sequencing at a depth of 80 million reads per sample. Samples were sequenced using QIAseq miRNA preparation kit (QIAGEN, Netherlands) and the Illumina NovaSeq 6000 sequencing system (Illumina Inc., USA).

The resultant fastq file underwent quality control analysis using FastQC. Cutadapt was applied to remove the sequencing adaptor regions and filter out reads less than 15nt or greater than 25nt in length, and reads with unidentified base pairs. After applying FastQC for a second time, the reads were aligned to miRbase Version 21 (Kozomara et al., 2019) using Bowtie (Langmead, 2010), with Hg38 as the reference genome. The created Sequence Alignment Map (SAM) was converted into a Binary Alignment Map (BAM) file using HTSeq (Anders et al., 2015), which was used to create a raw count file. This process was repeated for each sample. RStudio was used to collate the raw reads and perform differential expression analysis with DESeq2 (Love et al., 2014). Data visualisation was achieved using ggplot2, pheatmap and patchwork

### Luciferase Assay

The PsiCheck-2-Pri-miR-17-92a was a kind gift from Professor Yu Zhou’s laboratory (Wuhan University, China). The cloning of the plasmid is described in their study on Primary microRNA (Pri-miRNA) processing (Jiang et al., 2017). The plasmid was received via post, dried on filter paper. The dried blots of plasmid DNA were eluted and underwent bacterial transformation. Plasmid DNA was isolated via QIAprep Spin Miniprep Kit (Qiagen, USA) or GeneJet Plasmid Miniprep Kit (Thermo Fisher, USA).

For the Dual-Luciferase assay, cells were seeded at a density of 2.5x10^4^ cells per well in a 24 well plate. At 60-70% confluency, cells were transfected with either the Psi-Check-2-WT PDCD4 (Ajuyah et al., 2019) or Psi-Check-2-pri-miR-17-92a plasmid in combination with 20pmol miR-21 mimic (Applied Biosystems^TM^, Thermo Fisher, Australia), Let-7a mimic (Applied Biosystems^TM^, Thermo Fisher, Australia), or scramble control. Plasmid transfection only and untransfected cells were included as transfection controls. After 24 hours, the Firefly and Renilla Luciferase activity was measured in triplicate for each sample using the Dual-Luciferase Reporter Assay System (Promega, Australia). Luciferase activity was expressed as a ratio of Renilla to Firefly activity (Van Etten et al., 2013).

### Network Analysis

Cytoscape version (Shannon et al., 2003) 3.3.0 was used to visualise the relationships between the miRNAs identified as dysregulated using the MicroRNA OpenArray, along with their target genes. MiRNA target information was sourced from miRTarBase (Chou et al., 2018), and the *Homo sapiens* BIOGRID (Stark et al., 2006) interactome was accessed via Cytoscape. The targets of each identified miRNA were isolated from the *H. sapiens* interactome and merged to create a miRNA:miRNA:gene network.

Gene targets were annotated as transcription factors (TF), oncogenes (ONC), or tumour suppressor genes (TSG) according to TransmiR (Wang et al., 2010), ONGene (Liu et al., 2017), and TSGene (Zhao et al., 2016). To filter the network, genes that were classified as all three annotations were selected with their direct neighbours and imported as a new interactome. The result of this analysis is the creation of a network that identifies the potential effect of miRNA:miRNA interactions on TF’s, ONC’s, and TSG’s. Network analysis was performed within Cytoscape to identify influential miRNAs and genes. Of interest are the values for Node Degree, meaning the number of edges connected to a node, and Betweenness Centrality, which is the degree to which a node influences a network.

### Bioinformatics

Binding site prediction was performed using the command line tools miRanda and RNAHybrid (Kruger and Rehmsmeier, 2006). miRbase (version 22) (Kozomara et al., 2019) was used to extract the mature miR-21-5p sequence (MIMAT0000076), whereas the MIR17HG sequence was sourced from the NCBI Nucleotide database (Reference Sequence: NR_027350.1).

### Statistical Analysis

Fold change values were compared between samples using an unpaired two-tailed student’s t-test in GraphPad Prism (Version 8.0.1), with a p-value <0.05 considered statistically significant. Expression values were visualised as mean ± standard deviation.

## Results

### The Identification of miRNAs influenced by miR-21 overexpression

To determine the consequence of miR-21 on endogenous miRNA expression, we conducted a TaqMan MicroRNA Open Array on UMSCC22B samples transfected with a miR-21 mimic and compared the resultant miRNA expression to untransfected cells (Supplementary Figure 1).

Fold change analysis and filtering by amplification efficiency determined that 10 miRNAs were upregulated, while 150 miRNAs were downregulated compared to their expression in the non-transfected control (Supplementary Table 1). The top 10 most upregulated and downregulated miRNAs and their fold changes are shown in Table 1.

**Table 1:**
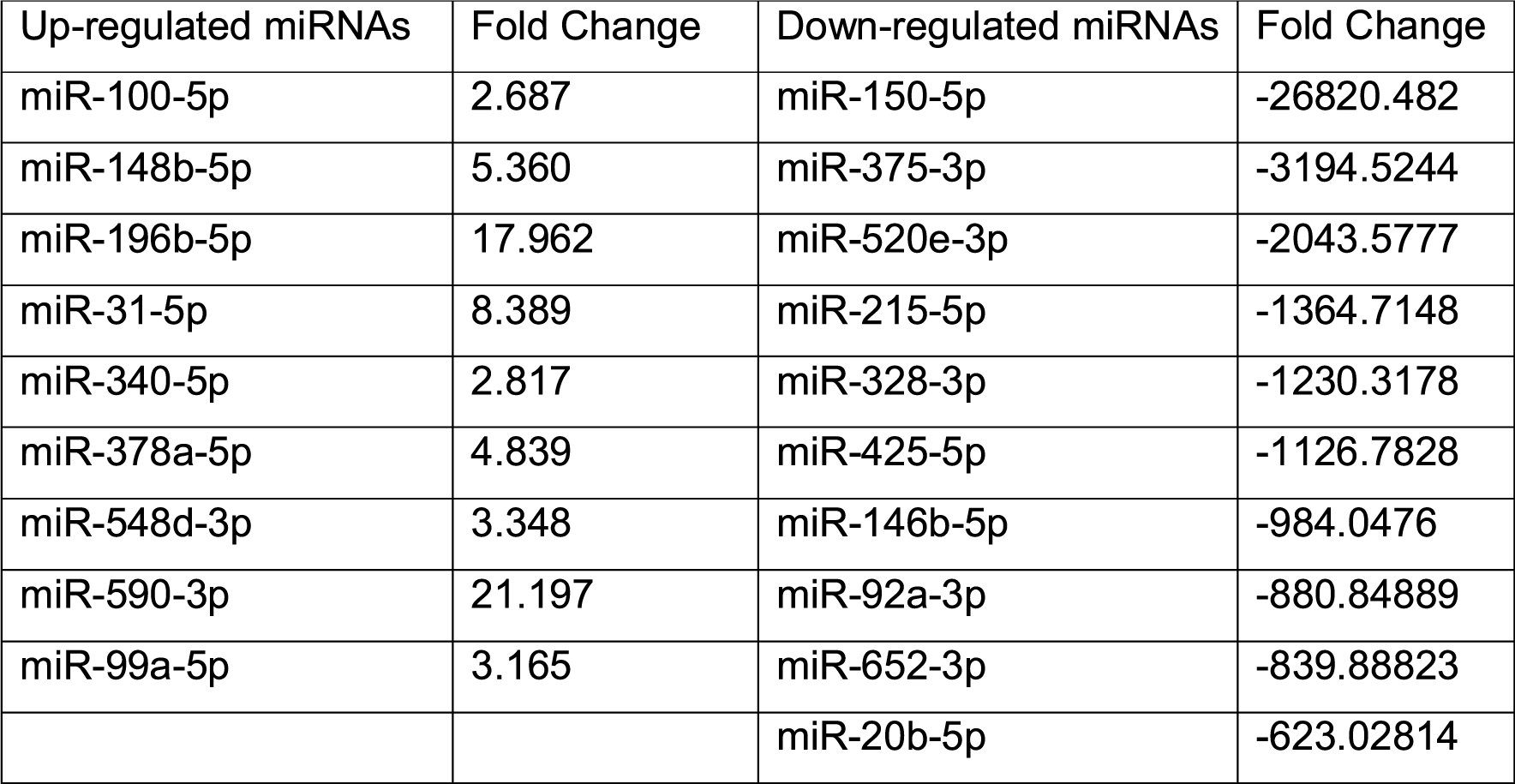
Curated list of dysregulated miRNAs in cancer cells overexpressing a miR- 21 mimic.

A series of miRNA:miRNA gene networks were created in Cytoscape from the information gathered from the OpenArray, with the aim of identifying potential pathways and mechanisms to explain the dysregulation of the identified miRNAs, and their potential role in cancer development. Firstly, the upregulated miRNAs in response to miR-21 were collated, and their targets were extracted from miRTarBase. An extensive network of all the listed miRNAs and genes was created from this information and contained 2383 nodes and 5232 edges. Filtering was applied to the complete network to identify common roles or potential pathways adjoining the miRNAs. This was performed by selecting for a particular gene annotation (ONC, TSG, or TF) and all direct neighbours of the selected genes.

This resulted in a series of networks filtered by the presence of ONC, TSG, or TF- annotated genes. Further filtering was applied whereby genes labelled as ONC, TSG, and TF inclusive, and their direct neighbours were selected to form a new network. Applying this filtering process to the network of upregulated miRNAs with miR-21 resulted in an interactome with 88 nodes and 339 edges (Figure 1A). As expected, the node for miR-21 is the largest, indicating that it has the greatest number of connections. Of the ten identified upregulated miRNAs with miR-21 overexpression, only five were present in this final filtered network. Clustering of the nodes within the network using ClusterMaker (Morris et al., 2011) showed four distinct groups, centred around miR-21, Transcription factor Sp1 (SP1), Tumour Protein p63 (TP63), SUZ12 Polycomb Repressive Complex 2 Subunit (SUZ12), and an axis between miR-340-5p, miR-548b-5p and miR-100-5p. Known targets for these miRNAs include known cancer genes such as Forkhead box protein O1 (FOXO1), Epidermal Growth Factor Receptor (EGFR), and Signal Transducer and activator of transcription 3 (STAT3).

**Figure 1.**
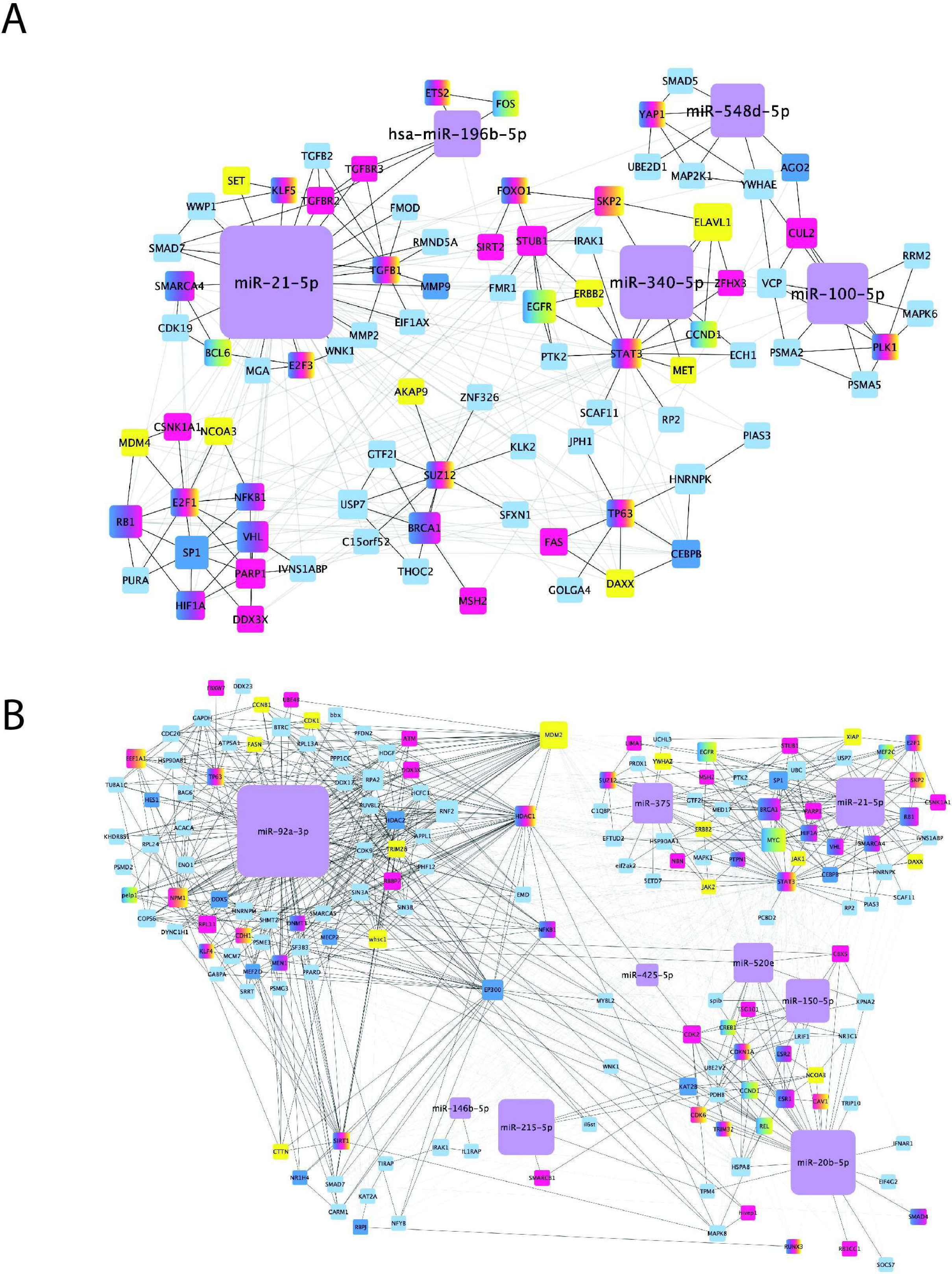
Gene:miRNA network integrating miR-21 and its Targets. In this network, miRNA, transcription factors (TF), tumour suppressor genes (TSG) and oncogenes (ONC) are represented with purple, blue, magenta and yellow, respectively. The size of the node is indicative of the number of connecting edges. Black edges indicate the main interactions as indicated by ClusterMaker, while the edges depicted in grey are background interactions. A) Gene:miRNA network integrating miR-21 and its upregulated miRNAs with their respective target genes, as filtered down by genes annotated with all three classifications (ONC, TSG, and TF), and their direct neighbours. B) A gene:miRNA network integrating miR-21 and its downregulated miRNAs with their respective target genes, as filtered down by genes annotated with all three classifications (ONC, TSG, and TF), and their direct neighbours

This process was repeated for the ten most downregulated miRNAs in response to miR-21 overexpression. The initial network had 4166 nodes and 14646 edges.

Filtering the network as before gave a resultant network of 320 nodes and 1980 edges (Figure 1B). When clusterMaker was applied, the nodes were placed into four distinct groups, with several miRNAs in each group. The major cluster in this network was formed surrounding miR-92a-3p, which connects to other clusters via Mouse double minute 2 homolog (MDM2), Histone Deacetylase 1 (HDAC1), and E1A Binding Protein P300 (EP300). Another node cluster connects miR-375 with cancer genes such as EGFR, Breast cancer type 1 susceptibility protein (BRCA1), and MYC Proto-Oncogene (MYC). The third cluster connects five miRNAs: miR-20b-5p, miR-150-5p, miR-520e, miR-215-5p, and miR-425-5p. The remaining small cluster predominantly consists of miR-146b-5p and its gene targets.

Several differences were observed in the networks of miRNAs that were either upregulated or downregulated by miR-21. The network associated with downregulated miRNAs exhibited a larger number of nodes and connections compared to the network for upregulated miRNAs. Additionally, there were fewer miRNAs present in the network of upregulated miRNAs when compared to the network of downregulated miRNAs. These observations suggest that the downregulation of miRNAs due to overexpression of miR-21 has a more significant impact than the effects observed with the upregulated miRNAs.

### miR-21 regulates the expression of the miR-17∼92a cluster members

We noted that miR-92a-3p levels were decreased upon miR-21 overexpression. This miRNA is part of the miR-17∼92a cluster, which is dysregulated in several cancers, including B cell lymphoma (Yan et al., 2019), carcinoma of the lung (Ventura et al., 2008), ovary (Deng et al., 2023), and colorectal cancers (Nishida et al., 2012). To demonstrate that miR-92a and cluster members (miR-17, miR-18, miR-19a, miR- 20a, and miR-19b-1) are potentially regulated by miR-21, UMSCC22B cells were transfected with the miR-21 mimic and a miR-21 ASO (antisense oligonucleotide).

For the miR-21 mimic and ASO, we recorded a significant increase and decrease in miR-21 expression levels, respectively (Figure 2A). We then assessed the expression levels of individual members of the miR-17-92a cluster using qRT-PCR within these transfected cells. We transfected miR-21 at two concentrations of 10 pmol and 30 pmol, with final expression levels calculated relative to the scramble control. In the miR-21 overexpressing cells, nearly all the cluster members displayed reduced expression levels with a greater decrease at 30 pmol of miR-21 (Figure 2B- F). In cells containing the miR-21 ASO, whereby miR-21 was depleted, there was a general trend in that the miRNAs in this cluster returned to baseline, similar to levels seen in the scramble control. This would suggest that the reduction in miR-21 mediates an increase in the observed levels for miR-17, miR-18, miR-19b, miR-20 and miR-92a.

**Figure 2:**
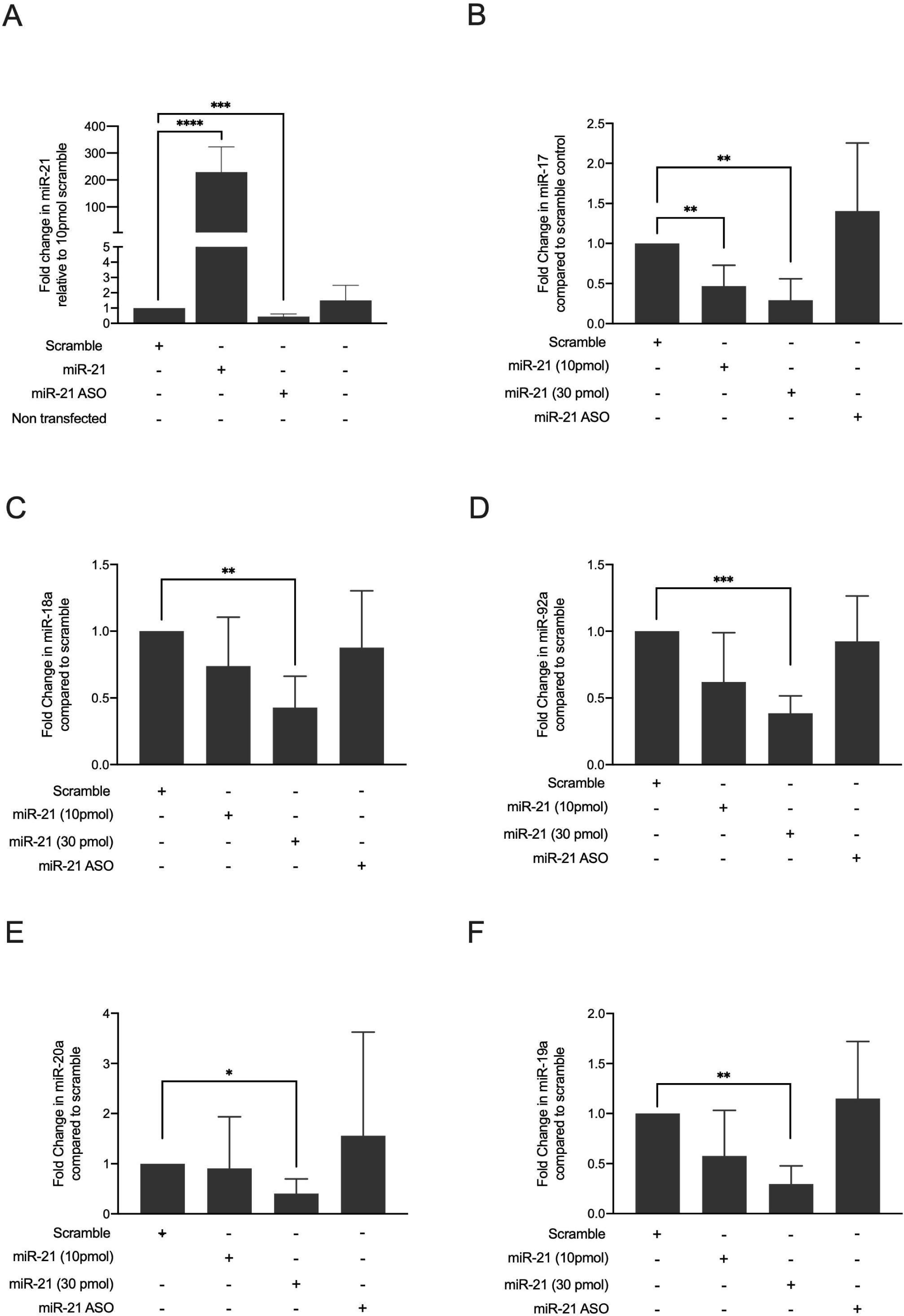
Expression levels for miR-17-92a cluster members in UMSCC22B cells transfected with miR-21 mimic (10pmol and 30pmol) and ASO normalised to the scramble control. A) Levels of miR-21 in cells transfected with the miR-21 mimic and ASO. B) Levels of miR-17. C) Levels of miR-18a. D) Levels of miR-92a. E) Levels of miR-20a. F) Levels of miR-19a.

To consolidate our initial observation, the total RNA from these transfected cells were sent for small RNA sequencing. After read normalisation, the expression levels of the miR-17∼92a cluster members were examined in relation to the addition of miR-21 or miR-21 ASO (Figure 3). All miRNA cluster members exhibited the same trend as observed in the qRT-PCR analysis. The presence of miR-21 led to a decrease in expression, while blocking of miR-21 resulted in elevated expression levels across all cluster members. Notably, cells treated with miR-21 ASO showed significantly higher miRNA expression levels than those treated with the scramble control. To determine if this pattern between miR-21 and the cluster was observed in a clinical context, we extracted miRNA expression data for nine common cancers from the TCGA database (Supplementary Figure 2). In all instances, miR-21 was highly expressed, whereas members of the cluster were noted at lower expression levels. This was very consistent across all nine primary cancers. These results support the idea that miR-21 could potentially regulate individual members of the miR-17-92a cluster.

**Figure 3:**
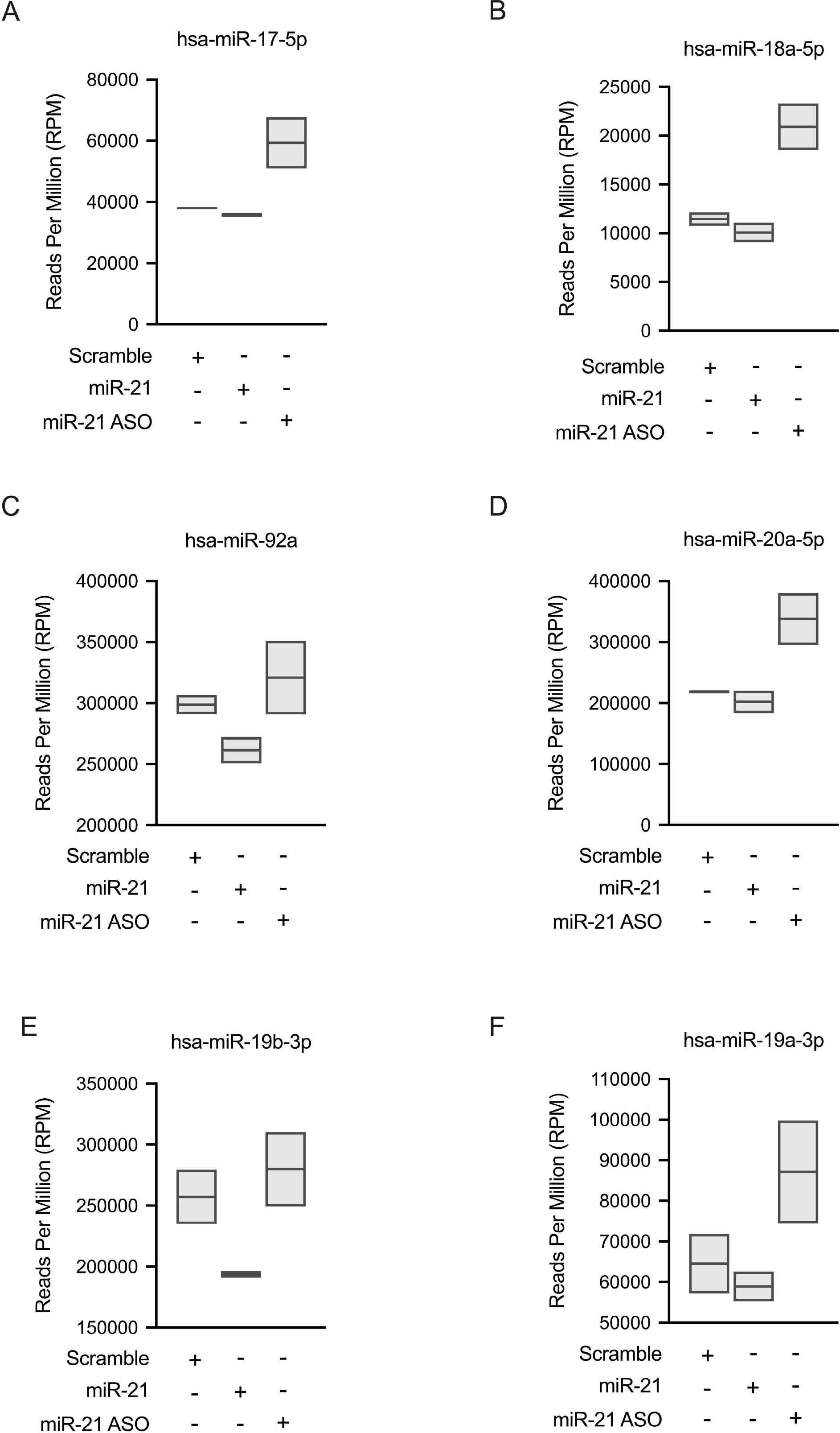
Small RNA sequencing of the UMSCC22B cells transfected with the miR- 21 mice and ASO. The box plots represent the normalised reads for each member of the miR-17-92a cluster derived from triplicate transfections for each treatment.

### The miR-17∼92a cluster harbours miR-21 binding sites

Our findings prompted further exploration into the mechanisms underlying the interaction between miR-21 and the miR-17-92a cluster. Other studies have highlighted the direct binding of miRNAs to MREs on target primary miRNAs as a direct mechanism. Following this line of inquiry, we analysed MIR17HG, which is the host gene for the miR-17∼92a cluster, for potential miR-21 binding sites. Utilising miRanda and RNAhybrid, we identified several miR-21 binding sites within MIR17HG. The miRanda tool mapped a specific site between positions 2385 and 2405 with a binding energy of -15.16 kcal/mol, indicating a potential binding of miR- 21’s seed sequence at positions 2-8 and additional binding at the 3’ end (Supplementary Table 2). Furthermore, RNAhybrid identified five other miR-21 binding sites across the 5000bp length of MIR17HG, with binding energies between - 25 and -20 kcal/mol, (Supplementary Table 3).

Using this data, we created a graphical representation to illustrate the predicted miR- 21 binding sites within MIR17HG, placing them in context with the cluster members (Supplementary Figure 3). This analysis revealed that only one of the predicted miR-21 sites, identified by RNAhybrid, located from positions 1663 to 1690 of the MIR17HG transcript, falls within the pre-miRNA region for the miR-17∼92a cluster. This site overlaps with the regions for miR-19b-1-5p and miR-19b-1-3p, suggesting a plausible direct interaction between miR-21 and MIR17HG.

### miR-21 can bind to the MIR17HG and regulate expression

To demonstrate that miR-21 can directly bind to MIR17HG, we employed the use of an in-vitro dual-luciferase reporter system. The coding region of pri-miR-17∼92a was inserted downstream of the luciferase gene. If binding occurs between miR-21 and the miR-17∼92a, a reduction will be observed in the relative luciferase activity.

Furthermore, a plasmid containing the 3’UTR of PDCD4 was used as a positive control to confirm miR-21 binding and activity (Ajuyah et al., 2019). These reporters were then transfected into UMSCC22B or HeLa cells with either the miR-21 mimic or the scramble control.

In the UMSCC22B cells, for the positive control harbouring the PDCD4 3’UTR, which contains a single known miR-21 MRE, we observed the expected decrease in relative luciferase units in the presence of miR-21. A similar reduction in activity was not seen in the scramble control (Figure 4A). When cells were then transfected with the Psi-Check-2-pri-miR-17-92a with miR-21, we noted a decrease in relative luciferase units, with no change observed with the scramble control (Figure 4B).

**Figure 4:**
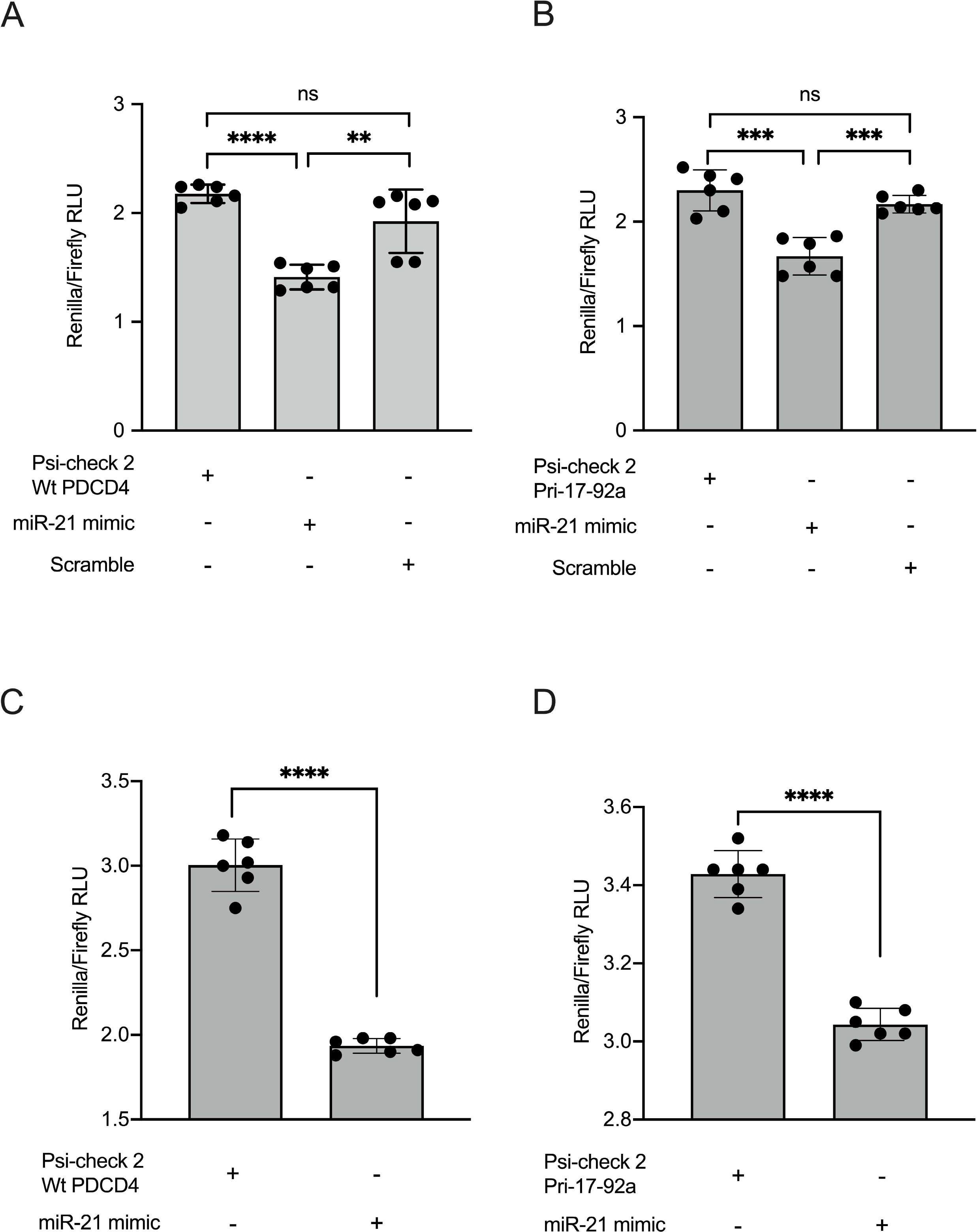
Luciferase assay to determine the direct binding of miR-21 to miR-17-92a. A) UMSCC22B cells transfected with a positive control Psi-Check 2 Wt PDCD4 plasmid in combination with miR-21 or scramble. B) UMSCC22B cells with the Psi- Check Pri-17-92a plasmid in combination with miR-21 or scramble. D) Hela cells transfected with the positive control Psi-Check 2 Wt PDCD4 in combination with miR-21 or scramble. E) HeLa cells with the Psi-Check Pri-17-92a plasmid in combination with miR-21 or scramble.

These constructs were also introduced into HeLa cells with similar outcomes in luciferase activity (Figure 4C-D). This overall decrease in luciferase units with the reporter Psi-Check-2-pri-miR-17-92a indicates that miR-21 can bind to pri-miR-17- 92a, leading to a suppression of luciferase expression.

There is evidence that some miRNAs can bind to specific primary miRNA transcripts and induce silencing at the post-transcriptional stage (Wang et al., 2014). To investigate whether miR-21 exhibits this behaviour, we measured the primary miR- 17-92a mRNA levels in cells that had been transfected with miR-21 (Figure 5). In the UMSCC22B cells, there was a significant reduction in the levels of primary miR-17- 92a mRNA when miR-21 was overexpressed. In comparison, cells containing the scramble mimic did not have a similar decrease (Figure 5A).

**Figure 5:**
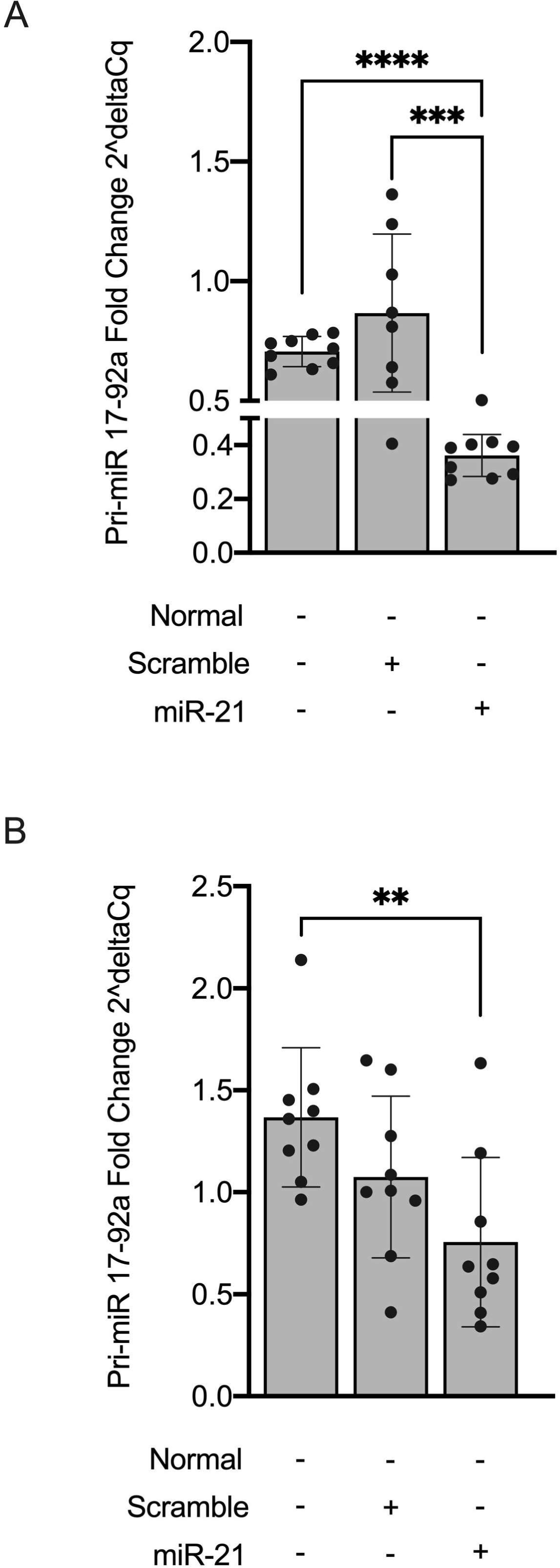
Reduction in Pri-miR-17-92a RNA in cells transfected with miR-21 and scramble A) 293 cells B) UMSCC22B cells.

In HeLa cells, a similar trend was observed, albeit not to the same levels. We do, however see that pri-miR-17-92a mRNA was significantly reduced when compared to untransfected cells (Figure 5B). It is noted that the scramble control also affects the levels of pri-miR-17-92a. Given these results, it is possible that miR-21 can bind to its MRE within the miR-17-92a cluster, thereby impacting expression levels.

### miRNAs Dysregulated by miR-21 can affect specific pathways

Gene Ontology (GO) analysis was performed for the networks of the miRNAs upregulated and downregulated by miR-21. This was conducted within Cytoscape using the plug-in BiNGO (Maere et al., 2005). The list of GO terms were then converted to their equivalent ID and input into REVIGO (Supek et al., 2011) to conduct semantic similarity analysis. The semantic similarity was visualised as a series of plots within R studio (Figure 6A). The top-most GO terms for the genes associated with the upregulated and downregulated miRNAs are also included in Supplementary Table Supplementary Tables 4 and 5, respectively.

**Figure 6:**
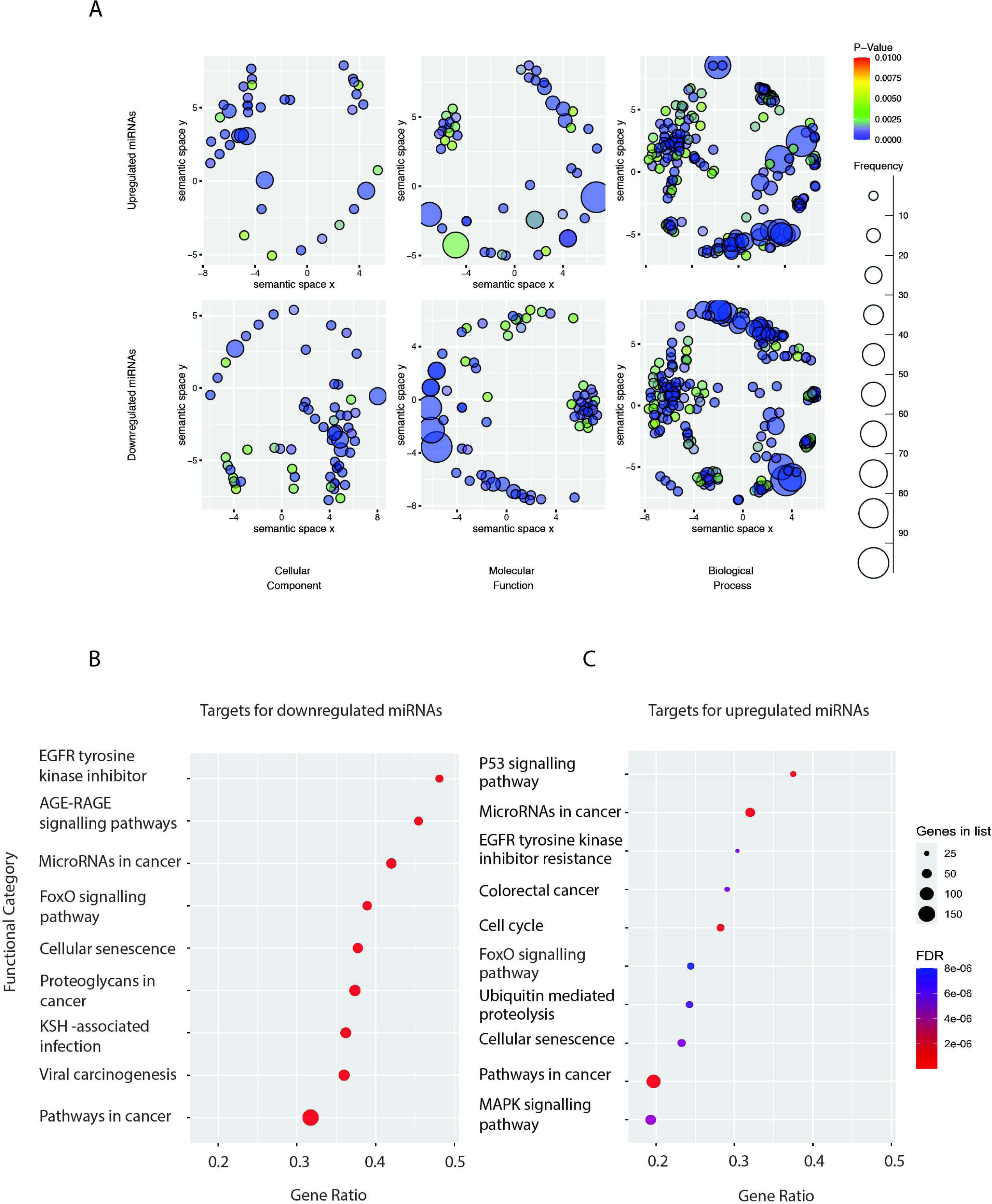
Summary of the GO terms identified for the miRNAs upregulated and downregulated with miR-21, separated by the GO classifications Cellular Component (A and D), Molecular Function (B and E) and Biological Process (C and F). For each graph, the size indicates the frequency of the GO term, and the colour scale is representative of p-value. The x and y axes measure the semantic similarity of the GO terms for the specific GO classification using the SimRel algorithm. G) KEGG terms plotted as gene ratio for upregulated miRNAs, H) KEGG terms plotted as gene ratio for downregulated miRNAs.

Of the GO categories, both the upregulated and downregulated miRNAs were the most involved in biological processes, as indicated by the high number of GO terms in this category and their associated p-value. From further observations, the downregulated miRNAs were also strongly involved in molecular function, as shown by the high frequency of GO terms and their significant p-value. It is unsurprising that the miRNAs influenced by miR-21 are extensively involved in biological processes.

KEGG analysis was performed for each of the miRNA:miRNA:gene networks through the ShinyGO database (Ge et al., 2020). The top ten most significant KEGG pathways identified for the miR-21 mediated upregulated and downregulated networks were visualised as a dot plot (Figures 6B and 6C respectively, or Supplementary Table 6)

The genes associated with the miR-21 upregulated miRNAs had KEGG terms associated with cancer development. Specific pathways identified included ’Cell cycle’, ’EGFR tyrosine kinase inhibitor resistance’, ’Cellular senescence’, ’Ubiquitin mediated proteolysis’, and ’FoxO signaling pathway’. Together with the additional identified pathways, this may suggest a significant contribution of these genes to the observed alterations in cancer development. Visualisation of the genes involved in the term ‘Pathways in Cancer’ are shown in Supplementary Figure 4

On the other hand, the analysis of the network formed by the downregulated miRNAs and their target genes revealed a different set of KEGG pathways, also closely tied to cancer progression but with additional links to viral infections. This network was particularly associated with pathways like ’Proteoglycans in cancer’, ’Kaposi’s sarcoma-associated herpesvirus infection’, and ’Hepatitis B’. The genes linked to the ’Pathways in Cancer’ KEGG annotation are shown in Supplementary Figure 5. The KEGG analysis of the two miR-21 mediated networks indicated that these genes may be highly involved in cancer processes.

## Discussion

MicroRNAs function as gene regulators by binding to nascent mRNA targets and are often dysregulated in cancer cells. There is an emerging idea that miRNAs can regulate other miRNAs at various junctions in the biogenesis pathway (Hill and Tran, 2021a). However, our understanding of how these regulatory interactions occur and which specific miRNAs are involved remains limited. To further contribute to this area, we sought to understand if miR-21, one of the most ubiquitously overexpressed miRNAs in cancer cells, could impact the expression levels of other miRNAs and to demonstrate specific examples of miR-21-mediated regulation.

In our *in vitro* model, it was observed that more miRNAs were downregulated in response to miR-21 transfection compared to those that were upregulated. This fits with the model that overexpressing miR-21 can directly bind to regions within the target miRNA to stall its biogenesis or affect processing at a specific stage. For the observed upregulation in miRNAs in response to miR-21 overexpression, it may be that miR-21 targets transcriptional inhibitory factors, and that the introduction of miR- 21 reduces these, resulting in the observed increases in specific miRNAs.

Additionally, there is evidence to suggest that miRNAs might increase their own transcription by directly binding to their targets (Orom et al., 2008).

The impact of the interactions between miR-21 and other miRNAs was further assessed by using a combination of miRNA:target tools to create a miRNA:RNA interactome. We see that the majority of targets are transcription factors for both the down or up-regulated miRNAs. From this mapping, two miRNAs, miR-100-5p and miR-92a-3p, stood out as the most influential due to the number of connections in each network. Consistent with our results, miR-100-5p has been shown to be upregulated in Head and Neck cancers (Wu et al., 2014). This miRNA is also involved in the Transforming Growth Factor Beta (TGFβ), Mitogen-activated Protein Kinase (MAPK), and Tumour Protein p53 (p53) cancer pathways (Chen et al., 2018) as identified in our KEGG analysis. Furthermore, many of the miR-100-5p targets are, in fact, RNA-binding proteins. The possible association between miR-21 and miR-100-5p could, therefore, drive transformational changes in some cancer cells.

Of the downregulated miRNAs, miR-92a was the most influential within the mapped network. This sparked our interest as miR-92a is part of the miR-17∼92a cluster, which encodes for six mature miRNAs (Olive et al., 2009) and members of this cluster are often dysregulated in several different cancers (Huang et al., 2021; Kim et al., 2012; Raponi et al., 2009). When the miR-21 mimic was introduced, we observed a notable reduction in the expression levels of all miR-17∼92a cluster members, suggesting that miR-21 could potentially regulate this cluster. Further evidence from the TCGA showed elevated levels of miR-21 with concomitantly reduced levels of the cluster. Given these findings, there are several different mechanisms that could account for the changes in the miR-17∼92a cluster in response to miR-21.

One of the potential mechanisms of regulation is the direct binding of miR-21 to pri- miR-17∼92a, thus inhibiting its production. Six predicted miR-21 binding sites were mapped to pri-miR-17∼92a. One binding site was located between the precursor sequences for miR-19b and miR-92a whereas the other sites were found outside the miR-17∼92a precursor sequence. Both the reduced luciferase activity of Psi-Check- 2-pri-miR-17-92a and the observed decrease in the primary transcript in response to miR-21 overexpression suggest the direct binding of miR-21 to pri-miR-17∼92a.

There have been very few examples of miRNAs directly binding to precursor or primary miRNA sequences. The miR-21 primary strand itself is regulated by nuclear miR-122, which results in a decrease of mature miR-21 levels (Wang et al., 2018). Another example is miR-361, which can directly bind to the primary miR-484 to prevent its processing by Drosha (Wang et al., 2014). Given that every member of the miR-17-92a was reduced, we reason that miR-21 can interact with its MRE within the miR-17-92a cluster. This suggests that regulation occurs within the nucleus, and that either miR-21 is transported into the nucleus or there is a reservoir of miR-21 already present. We know from previous studies that miR-21 is found both in the nucleus and cytoplasm (Meister et al., 2004), and there is a growing compendium of nuclear miRNAs present within mammalian cells (Hu et al., 2023). Our findings differ from the other two examples in that we did not find an accumulation of the primary transcript. Instead, we noted a decrease in the expression levels of the primary target. Measuring the precursor using the iLIME approach (Le et al., 2022) would add further evidence that regulation is occurring at the primary level.

Although we present in vitro evidence of a possible interaction, it is also plausible that the miR-17∼92a cluster might encounter additional regulatory pressures from miR-21. As other miRNA:miRNA interactions have demonstrated, transcriptional regulators are often key colluders in the regulation occurring between miRNAs (Borzi et al., 2017). There are a number of miR-21 targets that play a role in the transcription of miR-17∼92a, including MYC, STAT3, and transcription factor Sp1 (SP1). SP1 is a known miR-21 target (Yang et al., 2012) and has been shown to bind to the promoter of the MIR17HG to promote its expression (Meng et al., 2020). MYC and STAT3 are both promoters of the miR-17∼92a cluster, as well as targets of miR-17 and miR-20a in autoregulatory loops (Sylvestre et al., 2007). The two transcriptional promoters are also known targets of miR-21 (Aguda et al., 2008; Shishodia et al., 2014). These three transcriptional regulators could be additional mechanisms by which miR-21 mediates the miR-17∼92a cluster.

The GO and KEGG analysis conducted on the miRNA:RNA interactomes revealed that many of the targeted genes have significant roles in cellular metabolism and cancer-related processes. The cell lines used in our study were Head and Neck Squamous Cell Carcinoma (HNSCC) cells, and in this context and we observed that p53 was overrepresented in these networks, which is consistent with reports that p53 is dysregulated in HNSCC cells, (Manikandan et al., 2015). The observed loss of p53 in HNSCC might be linked to changes within the miRNA landscape, suggesting that the disruption of miRNA regulation may contribute to p53 dysfunction.

Since p53 itself also regulates specific miRNAs (Bommer et al., 2007; Garibaldi et al., 2016; Xi et al., 2006), alterations in its expression could further disturb the miRNA environment, potentially exacerbating cancer progression. Additionally, other cancer pathways identified, such as the MAPK signalling and FoxO pathways, are deeply involved in the migration, proliferation, differentiation, and apoptosis of cancer cells (Hornsveld et al., 2018). These pathways are listed under various cancer types in KEGG terms, suggesting that the miRNAs and genes within our networks are common across these cancers, indicating shared pathological pathways.

A consistent finding across the networks was the term ’EGFR tyrosine kinase inhibitor resistance’. EGFR inhibitors are frequently used to treat HNSCC, but they can lead to resistance and inflammation (Abu-Humaidan et al., 2020). Notably, several miRNAs, including miR-21, have been linked to chemoresistance (Sheng et al., 2022). The interactions between miRNA and mRNA identified in our analysis could contribute to this resistance.

Our research has shed light on the expanded role of miR-21 in the regulation of miRNA networks within cancer cells, particularly emphasising its influence over the miR-17∼92a cluster. This study has provided another example and deepens our understanding of miRNA:miRNA interactions. The finding that miR-21 can both upregulate and downregulate miRNAs reveals a nuanced regulatory mechanism. We have provided a list of potential miR-21 miRNA targets, and this alone suggests that miR-21’s regulatory scope extends beyond direct mRNA target suppression, implicating it in broader gene regulatory networks to now include other microRNAs.

Further research should aim to dissect the specific mechanisms through which miR- 21 interacts with other miRNAs in the nucleus and explore how these interactions contribute to gene silencing, as well as the involvement of Ago2. The overarching mechanism is likely a combination of both direct and indirect regulation activated at different stages of the cell cycle or development.

The interactions noted between miR-21 and various other miRNAs, and their subsequent effects on downstream mRNA targets, suggest a broader regulatory matrix that plays a crucial role in cellular function and pathology. This complex network of miRNA interactions suggests that miR-21 does not operate in isolation but rather functions as part of a larger miRNA regulatory landscape. These miRNAs could either synergise or antagonise each other’s effects, leading to the fine-tuning of gene expression that is critical for maintaining cellular homeostasis. Deciphering these mechanisms remains challenging and will be crucial for advancing our understanding of miRNA roles in gene regulation and their broader implications in cancer and other diseases.

